# The Arabidopsis D27-like1 is a *cis*/*cis*/*trans*-β-carotene Isomerase that Contributes to Strigolactone Biosynthesis and Negatively Impacts Abscisic Acid Level

**DOI:** 10.1101/2022.06.07.495147

**Authors:** Yu Yang, Haneen Abuauf, Shanshan Song, Jian You Wang, Yagiz Alagoz, Juan C. Moreno, Jianing Mi, Abdugaffor Ablazov, Muhammad Jamil, Shawkat Ali, Xiongjie Zheng, Aparna Balakrishna, Ikram Blilou, Salim Al-Babili

## Abstract

The enzyme DWARF27 (D27) catalyzes the reversible isomerization of all-*trans*- into 9-*cis*-β-carotene, initiating strigolactone (SL) biosynthesis. Genomes of higher plants encode two D27-homologs, D27-like1 and -like2, with unknown functions. Here, we investigated the enzymatic activity and biological function of the Arabidopsis D27-like1. *In vitro* enzymatic assays and Expression in *Synechocystis* sp. PCC6803 revealed a yet not reported 13-*cis*/15-*cis*/9-*cis*- and a 9-*cis*/all-*trans*-β-carotene isomerization. Although disruption of *AtD27-like1* did not cause SL deficiency phenotypes, overexpression of *AtD27-like1* in the *Atd27* mutant restored the more-branching phenotype, indicating a contribution of *AtD27-like1* to SL biosynthesis. Accordingly, generated *Atd27 Atd27like1* double mutants showed more pronounced branching phenotype, compared to *Atd27.* The contribution of AtD27-like1 to SL biosynthesis is likely due to its formation of 9-*cis*-β-carotene that was present at higher levels in *AtD27-like1* overexpressing lines. In contrast, *AtD27-like1* expression correlated negatively with the content of 9-*cis*-violaxanthin, a precursor of abscisic acid (ABA), in shoots. Consistently, ABA levels were higher in shoots and also in dry seeds of the *Atd27like1* and *Atd27 Atd27like1* mutants. Transgenic lines expressing β-glucuronidase (GUS) driven by the *AtD27LIKE1* promoter and transcript analysis performed with hormone-treated Arabidopsis seedlings unraveled that *AtD27LIKE1* is expressed in different tissues and regulated ABA and auxin. Taken together, our work revealed a *cis*/*cis-*β-carotene isomerase activity that affects the content of both *cis*-carotenoid derived plant hormones ABA and SLs.

## INTRODUCTION

Carotenoids are versatile pigments, which are essential for plant survival due to their function in photosynthesis and photoprotection (Demmig-Adams, 1990). The carotenoid backbone consists of an extended, conjugated system of double bonds susceptible to oxygen attack yielding cleavage products called apocarotenoids, which act as small signaling molecules regulating growth and development, stress response, and the aroma and color of flowers and fruits (D’Alessandro et al., 2018; Jia et al., 2019; Wang et al., 2019; Shi et al., 2020). Additionally, the breakdown of carotenoid generates the precursors of the phytohormones abscisic acid (ABA) and strigolactones (SLs) (Moreno et al., 2021; Zheng et al., 2021). ABA regulates several crucial processes in plants, such as seed dormancy and germination (Shu et al., 2016; Wang et al., 2020), shoot and root growth and development (Finkelstein, 2013; Cui et al., 2016; Gao et al., 2016; Diretto et al., 2020), stomatal closure and movement (Merilo et al., 2015), and is best known for mediating biotic and abiotic stress (Nambara and Marion-Poll, 2005; Chen et al., 2020). Indeed, ABA accumulates in response to drought, salt, and cold stress; enhancing tolerance to unfavorable conditions (Zeevaart and Creelman, 1988; Yamaguchi-Shinozaki and Shinozaki, 2006). ABA biosynthesis is well-characterized in different plant species (Nambara and Marion-Poll, 2005; Cutler et al., 2010; Dong et al., 2015), starting with the hydroxylation of β-carotene into zeaxanthin (Kim and DellaPenna, 2006) and followed by the double epoxidation of zeaxanthin into violaxanthin (Bouvier et al., 1996; Marin et al., 1996) that can be converted into neoxanthin (North et al., 2007). The subsequent isomerization of all-*trans*-violaxanthin/neoxanthin yields the corresponding 9-*cis*/9’-*cis*-isomer, respectively (Isaacson et al., 2002; Tan et al., 2003; Perreau et al., 2020). The cleavage of these two *cis*-epoxy-xanthophylls (C_40_) produces the ABA precursor xanthoxin (C_15_), representing the first committed and rate-limiting-step in ABA biosynthesis, which takes place in plastids and is mediated by 9-*cis*-epoxycarotenoid cleavage dioxygenase (NCED) enzymes (Qin and Zeevaart, 1999). Thereafter, ABA DEFICIENT 2 (ABA2) enzyme catalyzes the dehydrogenation of xanthoxin to abscisic aldehyde in the cytosol, which is then oxidized to ABA by abscisic aldehyde oxidase (AAO) that requires a molybdenum cofactor encoded by *ABA3* (Schwartz et al., 1997). A recent study reported a non-canonical, zeaxanthin epoxidase (ABA1) independent, biosynthetic route in which β-apo-11-carotenoids serve as novel ABA precursors (Jia et al., 2022).

SLs were originally discovered as as host-derived germination stimulants for root parasitic plants, such as *Striga hermonthica* (Jamil et al., 2021). Later on, they were shown to be a mediator of arbuscular mycorrhizal symbiosis (). Moreover, SLs are an important plant hormone involved in various aspects of plant growth and development, such as inhibiting shoot branching (Gomez-Roldan et al., 2008) and shaping root architecture(Koltai, 2011). SL biosynthesis starts with the reversible isomerization of all-*trans*-β-carotene into 9-*cis*-β-carotene, which is catalyzed by the DWARF27 (D27) isomerase. In the next steps, 9-*cis*-β-carotene is converted by the CAROTENOID CLEAVAGE DYOXYGENASE7 and 8 (CCD7 and CCD8) into carlactone (CL), the precursor of SLs (Alder et al., 2012). The isomerase activity of D27 was unraveled through expression in engineered, β-carotene accumulating *E.coli* cells harboring the pBeta plasmid (Alder et al., 2012), which encodes the bacterial crtE, crtB, crtl and crtY enzymes catalyzing all-*trans-*β-carotene biosynthesis (Prado-Cabrero et al., 2007). *In vitro* assays performed with all-*trans*- and 9-*cis*-β-carotene confirmed that the isomerization is reversible with the reaction equilibrium preference for all-*trans-*β-carotene. Other *cis*-isomers, including 13- and 15-*cis*-β-carotene, were not isomerized, indicating the specificity of *At*D27 and *Os*D27 enzymes for the C_9_-C_10_ double bond (Bruno and Al-Babili, 2016; Abuauf et al., 2018). Besides, CCD7 enzymes also cleave hydroxylated, bicyclic 9-*cis*-configured carotenoids, such as 9-*cis*-zeaxanthin and 9-*cis*-lutein, leading to hydroxyl-carlactone that might be a precursor of yet unidentified SLs (Baz et al., 2018). Same as SLs, ABA biosynthesis requires 9-*cis*-configured carotenoids as precursors. This raises the question of whether D27 enzymes can produce 9-*cis*-xanthophylls. However, further *in vitro* assays indicated that D27 enzymes isomerize the C_9_-C_10_ double bond adjacent to unmodified β-ionone ring in carotenoid substrates, showing the isomerization of all-*trans*- or 9-*cis*-configured β-carotene, α-carotene, and cryptoxanthin *in vitro*, but not violaxanthin or neoxanthin (Bruno and Al-Babili, 2016; Abuauf et al., 2018). The inhibition of D27 isomerization by adding silver acetate suggests the presence of an iron cluster in the D27 reaction center (Harrison et al., 2015). *D27* encodes a plastid-localized enzyme mainly expressed in immature flowers in Arabidopsis and vascular cells of rice shoots and roots (Lin et al., 2009; Abuauf et al., 2018). The phenotype of rice and Arabidopsis *d27* mutants resembles that of other SL-deficient mutants (rice *d17* and *d10*, and Arabidopsis *max3* and *max4*) but is less pronounced than that of *ccd7* or *ccd8* mutants (Lin et al., 2009; Waters et al., 2012). This mild phenotype indicated that the loss of D27 activity might be compensated by photoisomerization and/or by the activity of D27 homolog(s), i.e., D27-LIKE1 and D27-LIKE2, which are conserved in land plants (Waters et al., 2012). However, the enzymatic activity and biological functions of the D27 homologs remained elusive.

In plants, SLs and ABA derive from the same precursor β-carotene. The rice SL-deficient *d10* and *d17* mutants accumulate higher amounts of shoot ABA than wild-type, especially under drought stress conditions. However, the rice *d27* mutant displays reduced drought tolerance and accumulates less leaf ABA content, indicating D27 might be a possible hub modulating the SL and ABA accumulation in plant tissues (Haider et al., 2018), which was supported by the decreased expression levels of the ABA-responsive genes *OsMYB2* and *RAB16C* in roots and shoots of the rice *d27*. Moreover, the *Osd27like1* and *Osd27like2* mutants showed reduced ABA levels in shoots, suggesting a possible contribution of the rice D27 family to ABA biosynthesis (Liu et al., 2020).

In this work, we set out to characterize the enzymatic activity of the Arabidopsis D27-like1 and to investigate its possible involvement in SL and ABA biosynthesis. For this purpose, we performed *in vitro* and *in vivo* activity tests, generated knock-out mutants and overexpressing lines, phenotyped them and determined their carotenoids and ABA content. Our results demonstrate that AtD27LIKE1 is a β-carotene isomerase that catalyzes yet not described *cis*/*cis*-isomerization reactions, in addition to 9-*cis*/all-*trans*-conversion reported for D27 enzymes, and that *AtD27LIKE1* is involved in the biosynthesis of the two carotenoid-derived plant hormones ABA and SLs, by contributing to SL biosynthesis while negatively impacting ABA content.

## RESULTS

### *At*D27LIKE1 catalyzed novel *cis* to *cis* β-carotene isomerization reactions

Carotenoid-accumulating *E. coli* strains are an efficient system for characterizing carotenoid-metabolizing enzymes. Therefore, we expressed the *At*D27LIKE1 protein fused to thioredoxin, encoded in pThio-*At*D27LIKE1, in β-carotene-, lycopene- and zeaxanthin-accumulating *E.coli* cells (Matthews and Wurtzel, 2000; Prado-Cabrero et al., 2007) and used thioredoxin-*At*D27 as a comparator. Introduction of *At*D27LIKE1 into β-carotene-accumulating *E. coli* cells did not change the pattern of β-carotene isomers, while *At*D27 caused the expected increase in the 9-*cis*/all-*trans*-β-carotene ratio (**Supplemental Figure S1A-B**). Similarly, we did not observe any isomerization activity upon introducing pThio-*At*D27LIKE1 in all-*trans*-lycopene- and zeaxanthin-accumulating *E. coli* cells (**Supplemental Figure S1C-D**). β-Carotene occurs naturally in 4 different stereo-configurations, i.e. in all-*trans*-, 9-*cis*-, 13-*cis*-, and 15-*cis*-configuration (for structures, see Figure 1). Because the *cis*-isomers are not accumulated in carotenoid-producing *E. coli* strains, we performed *in vitro* assays with different β-carotene geometric isomers, using crude lysates of thioredoxin-*At*D27LIKE1 and -*At*D27 expressing BL21 *E.coli* cells that are equipped with the pGro7 plasmid that encodes chaperones improving protein folding. Incubation of thioredoxin-*At*D27LIKE1 with all*-trans*-β-carotene did not change the content of any *cis* isomer (**Supplemental Table S2**), while thioredoxin-*At*D27 significantly increased the 9-*cis*/all-*trans*-β-carotene ratio (Figure 1A-B), which was in line with the *in vivo* assays (**Supplemental Figure S1A-B**). Incubation of thioredoxin-*At*D27 with 9-*cis*-β-carotene confirmed the previously reported conversion into the all-*trans*-isomer (Abuauf et al., 2018). Interestingly, *At*D27LIKE1 converted 9-*cis*-β-carotene in all*-trans*- and 13-*cis*-β-carotene (Figure 1B), increasing their proportions in total β-carotene content from 5% and 4% to around 40% and 15% (**Supplemental Table S3**), respectively. Thioredoxin-*At*D27LIKE1 also converted 13-*cis*-β-carotene into 9-*cis*-β-carotene, increasing the 9-*cis*/13-*cis* ratio from around 2% (**Supplemental Table S4**) in the control incubation to around 18 % (Figure 1C). We did not observe a conversion of 13-*cis*-β-carotene into the corresponding 15-*cis*- or the all-*trans*-isomer (Figure 1C). However, we detected the reverse reaction when incubating with 15-*cis*-β-carotene. As shown in (Figure 1D), the enzyme converted this substrate into 9-*cis*- and 13-*cis*-β-carotene, increasing their ratio from 1% and 9% to 7% and 13% (**Supplemental Table S5**), respectively (Figure 1D). As expected, we did not detect significant change in the isomers pattern upon incubating thioredoxin-*At*D27 with 13-*cis*- or 15-*cis*-β-carotene (Figure 1C-D). We also tested the activity of thioredoxin-*At*D27LIKE1 protein preparation on different carotenoids, i.e. all*-trans-*lutein, -violaxanthin, -neoxanthin, 9’*-cis*-neoxanthin and α-carotene. However, we did not observe any any isomerization activity (**Supplemental Figure S2A-E**), while thioredoxin-*At*D27 produced 9’-*cis*-α-carotene from the corresponding all-*trans*-isomer (**Supplemental Figure S2E-F**). These results suggest that *At*D27LIKE1 is a β-carotene isomerase that catalyzes yet not reported *cis*-*cis* isomerization, in addition to the conversion of 9-*cis*-into all-*trans*-β-carotene, which is known for Arabidopsis and rice D27 enzymes (Bruno and Al-Babili, 2016; Abuauf et al., 2018). Importantly, the enzyme formed 9-*cis*-β-carotene from other *cis*-configured β-carotene isomers, which indicated a possible contribution to SL biosynthesis.

**Figure 1.**
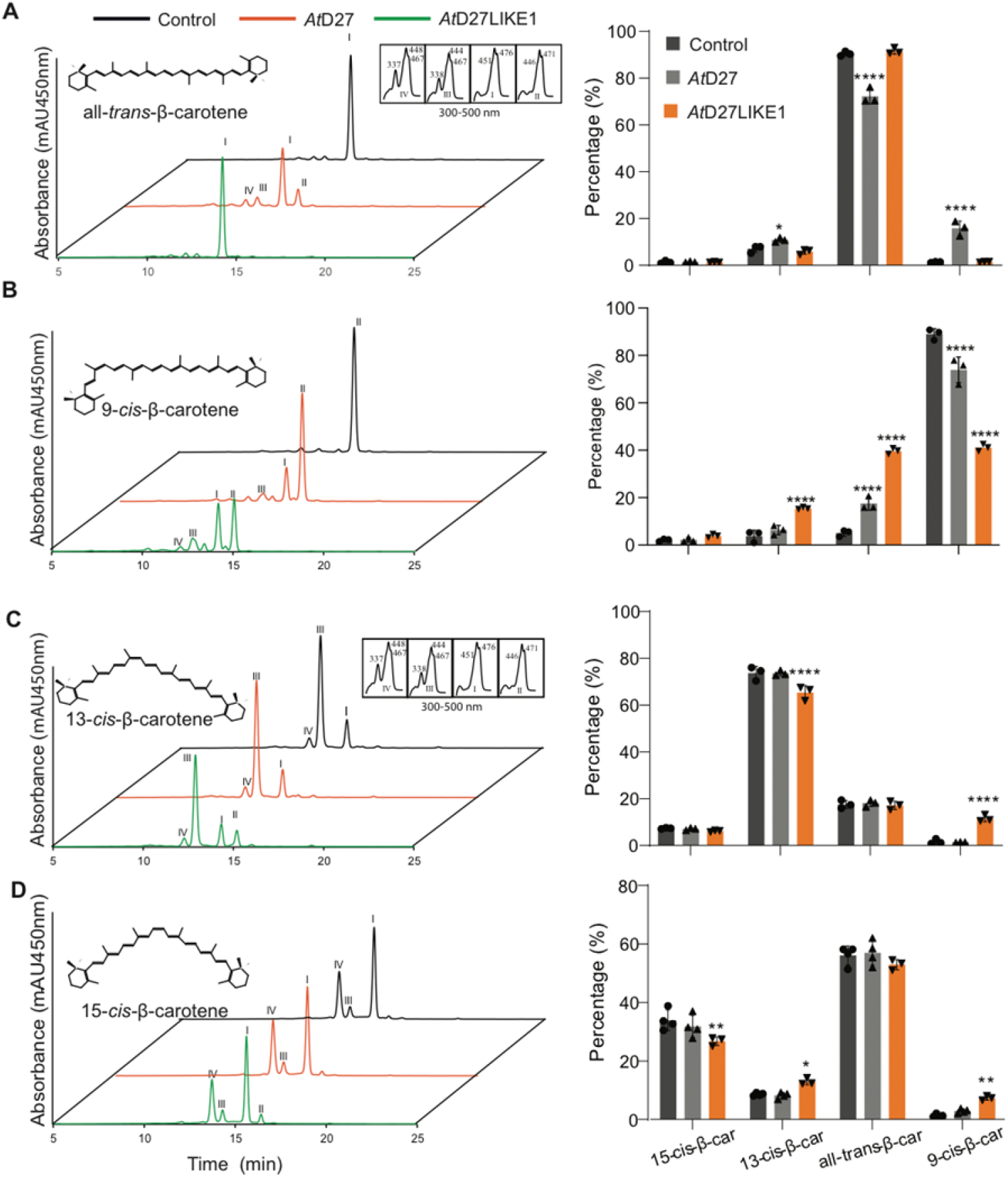
UHPLC analysis of *in vitro* assays performed with crude lysates of BL21 *E. coli* cells expressing thioredoxin-*At*D27 (AtD27), thioredoxin*-At*D27LIKE1 or thioredoxin (Control) with different β-carotene isomers. **A.** all-*trans*-β-carotene (peak I); **B.** 9-*cis*-β-carotene (peak II); **C.** 13-*cis*-β-carotene (peak III); **D.** 15-*cis*-β-carotene (peak IV). Left: Chromatograms of the incubations with different β-carotene isomers. **Right**: The relative peak surface of the different β-carotene isomers separated in the chromatograms (**Left**). Sum of all β-carotene peaks is considered as 100%. UV-Vis spectra are depicted in the insets. A non-paired two-tailed Student’s *t*-test was performed to determine significance (*n* = 3). *: p < 0.05, **: p < 0.01, ****: p < 0.0001. Error bars represent ±SD.

The occurrence of the *At*D27LIKE1 substrate β-carotene in plastids and the predicted presence of a plastid transit peptide indicate that *At*D27LIKE1 is a plastid enzyme. Therefore, we investigated the activity of *At*D27LIKE1 in the cyanobacterium strain *Synechocystis* sp. PCC 6803 (hereafter *Synechocystis*), considering the cyanobacterial origin of plastids. We first generated a *Synechocystis ΔcrtO* mutant lacking the β-carotene ketolase activity, CrtO encoded by *slr0088*, which converts β-carotene into echinenone(Fernández-González et al., 1997)(**Supplemental Figure S3**), assuming that blocking the echinenone branch might increase β-carotene content and change the isomer pattern. UHPLC analysis showed that the *ΔcrtO* mutant contained a higher content of *cis*-β-carotenes, compared to the wild type strain (**Supplemental Figure S4**), which made it a suitable system for testing *At*D27LIKE1 activity. Hence, we overexpressed *AtD27LIKE1* and simultaneously deleted *crtO* by generating a DNA fragment that combines the expression cassettes of *AtD27LIKE1* equipped with a C-terminal *His-tag* and kanamycin resistance and using it to replace c*rtO* via homologous recombination. To ensure a high expression level, we employed the strong constitutive promoter P*cpc560* (Zhou et al., 2014). (Figure 2A). Because cyanobacterial cells contain multiple chromosome copies (Mann and Carr, 1974), we verified the complete replacement of *crtO* in wild type alleles by PCR using specific primers (**Supplemental Table S1**). The *crtO* gene copies were completely lost and replaced by the *AtD27LIKE1*-expressing operon in all the *At*D27L1-OX lines, as only fragments specific for the inserted cassette were detected by PCR (Figure 2B). Moreover, we confirmed the presence of the *At*D27LIKE1 protein in all three *At*D27L1-OX lines by Western Blot, using anti His-tag antibodies (Figure 2C). Next, we extracted chlorophyll-a and carotenoids from the different lines, quantified chlorophyll and total carotenoids photometrically, and characterized the carotenoid pattern by UHPLC using a calibrated all-*trans*-β-carotene standard curve (**Supplemental Document 1**). The *ΔcrtO* mutant and *At*D27L1-OX lines displayed similar levels of chlorophyll-a, total carotenoids and total carotenoid/chlorophyll-a ratios (**Supplemental Figure S5**), indicating that the overexpression of *AtD27LIKE1* did not affected photosynthetic activity. However, UHPLC analysis revealed that the *At*D27L1-OX lines contained significantly reduced contents of 13-*cis*- and, particularly, 15-*cis*-β-carotene, while only one line showed a lower level of all-*trans*-β-carotene, compared to *ΔcrtO.* In the *ΔcrtO* mutant, the concentrations of 15-*cis*-β-carotene and 13-*cis*-β-carotene were around 0.064 μg/mL and 0.11 μg/mL, and the amounts decreased in the *At*D27L1-OX lines about ∼30%, respectively (Figure 2D). We also detected a decrease in the myxoxanthophyll content (**Supplemental Figure S5**), which might be indirectly related to the changes in *cis*-β-carotenes.

**Figure 2.**
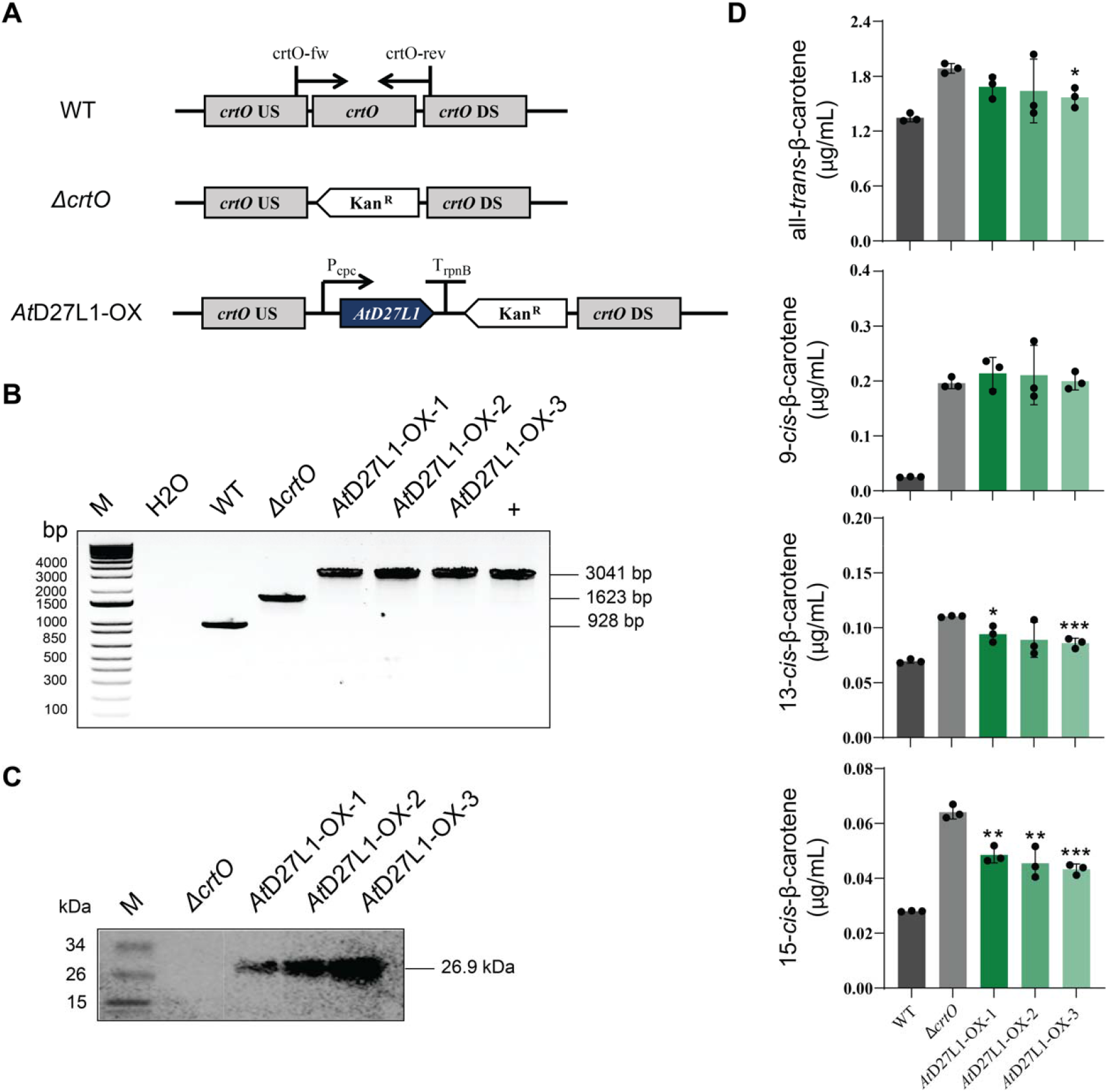
Characterization of transgenic *Synechocystis* sp. PCC 6803 and quantification of carotenoids. **A**. Schematic representation of the vectors for overexpression of Arabidopsis *AtD27L1* (*AtD27LIKE1*) by homologous recombination. **B.** Genotypic characterization of the *At*D27L1*-*OX strains. Complete segregation of mutant and overexpressing lines was verified by genomic PCR using primers crtO-fw/rev amplifying the whole insertion fragment as indicated. **C.** Western blot detection with protein extracts of *ΔcrtO* mutant and *At*D27L1-OX lines. The blot was treated with anti-His HRP conjugated antibody. The *At*D27L1 specific bands with a molecular weight of 26.9 kDa are indicated. **D**. Absolute quantification of β-carotene isomers in WT, *ΔcrtO* and *At*D27L1*-*OX lines. A non-paired two-tailed Student’s *t*-test was performed to determine significance (*n* = 3). *: p < 0.05, **: p < 0.01, ***: p < 0.001. Error bars represent ±SD. *At*D27L1: *At*D27LIKE1, *crtO* US/DS: upstream and downstream sequence of β-carotene ketolase gene of *Synechocystis* sp. PCC 6803: Kan^R^: kanamycin. Pcpc: promoter sequence. TrpnB: terminatior sequence. M: PAGERuler^TM^ Prestained Protein Ladder, 10-180kDa.

### Disruption and overexpression of *AtD27LIKE1* demonstrate its contribution to SL biosynthesis

To investigate the biological functions of *At*D27LIKE1 *in planta* and its possible contribution to SL biosynthesis, we generated *Atd27like1* and *Atd27 Atd27like1* knock-out mutants using CRISPR-Cas9 technology. For this purpose, we transformed *Arabidopsis* wild-type Col-0 and the previously described *Atd27* mutant (Waters et al., 2012) with the *At*D27LIKE1-CRISPR/Cas9 construct containing one gRNA that targets the first exon in *AtD27LIKE1* (Figure 3A). We obtained two independent Cas9-free homozygous *Atd27like1* mutant lines carrying each a single nucleotide insertion (T or A) in the target site of *AtD27LIKE1* at different positions, resulting in a frameshift and premature stop of *At*D27LIKE1. We also obtained two *Atd27 Atd27like1* double mutants, which carried either a 1-nucleotide insertion (T) or a 1-nucleotide insertion (A) and 36 bp deletion at a different position in exon-1 (Figure 3A) in the *Atd27* background. Next, we grew Col-0, *Atd27*, *Atd27like1*, *Atd27 d27like1* plants and recorded their height and axillary branching at 45-days-old stage. As shown in Figure 3B, we did not detect an increase in the number of shoot branches in the *Atd27like1* mutant, compared to the wild type; however, *Atd27* and *Atd27 Atd27like1* mutants showed a substantial increase of branches (Figure 3D), which was more pronounced in *Atd27 Atd27like1* than in the *Atd27* mutant. The *Atd27 Atd27like1* mutant showed also reduced plant-height (Figure 3C), a further phenotype caused by SL deficiency, which we did not observe in the *Atd27* mutant. These data indicate that *Atd27* and *Atd27like1* have overlapping function, pointing to a contribution of *AtD27LIKE1* to SL biosynthesis, likely by providing the 9-*cis*-β-carotene required for their biosynthesis. This contribution explains the more severe SL-deficiency phenotype of *Atd27 Atd27like1* double mutant, compared to *Atd27* mutant, and the relatively weak branching phenotype of *Atd27*, compared to *max3* or *max4* mutants (Waters et al., 2012).To determine the role of *AtD27LIKE1* in the biosynthesis of released SLs, we performed *Striga* seed germination assay with root exudates collected from hydroponically grown Col-0, *Atd27*, *Atd27like1*, and *Atd27 Atd27like1* plants, using the SL analog *rac*-GR24 (2.5 μM) as a positive control. Application of root exudates of Col-0 and *Atd27like1* plants resulted in the germination of around 10 % of the *Striga* seed, while root exudates from *Atd27* and *Atd27 Atd27like1-1* caused the germination of only ∼5 % (Figure 3E). As expected, application of GR24 showed much higher activity triggering the germination of ∼45 % of the seeds. These data demonstrated that the disruption of *AtD27LIKE1* did not affect the amounts of released SLs neither in wild-type nor in the *Atd27* background.

**Figure 3.**
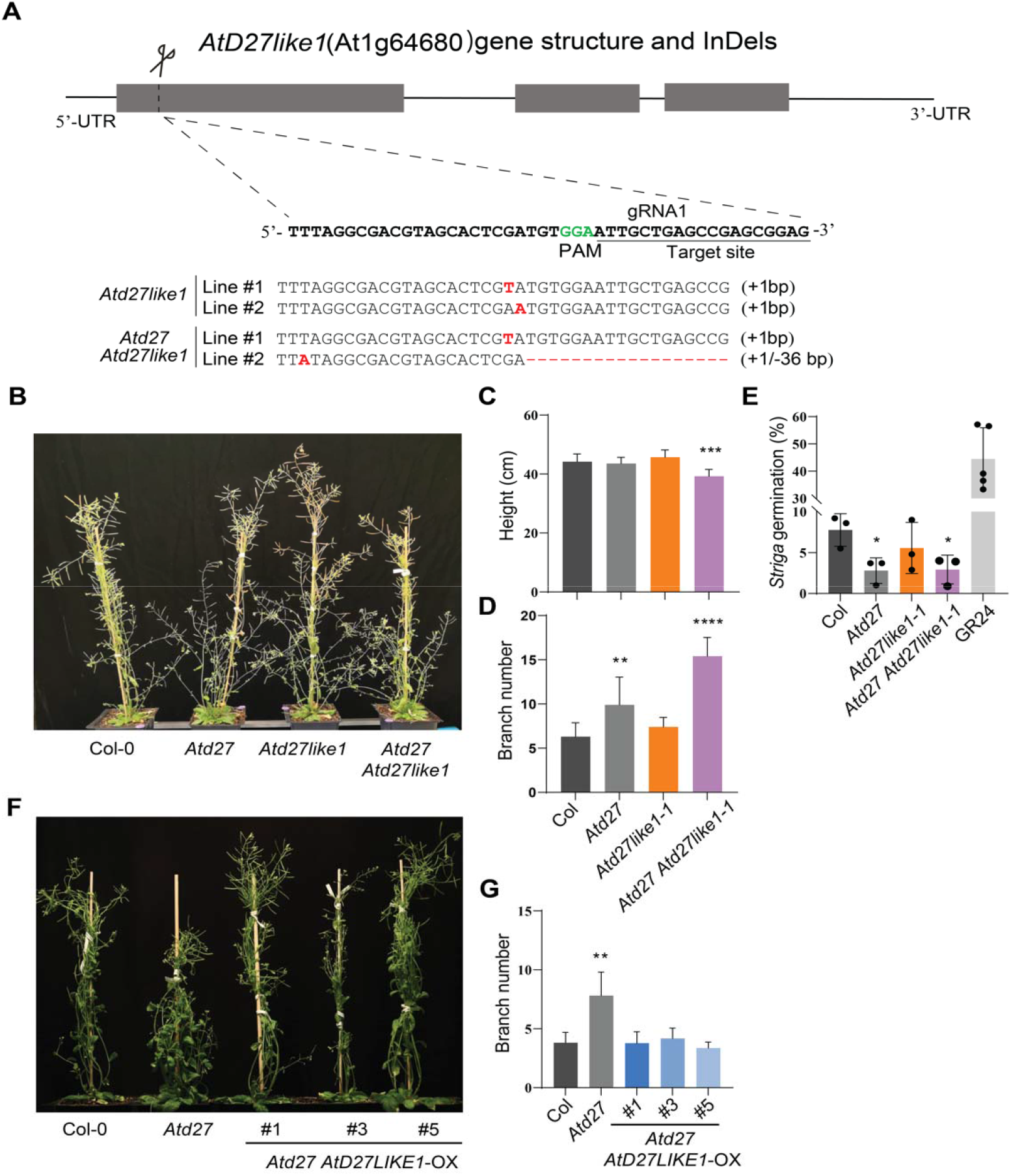
Generation and characterization of *Atd27like1*, *Atd27 Atd27like1*, and *Atd27 AtD27LIKE1-*OX. **A**. CRISPR Cas-9 protospacer adjacent motif (PAM)/guide RNA (gRNA) sequence and mutations of the *Atd27like1* and *Atd27 Atd27like1* CRISPR mutants. DNA insertions and deletions (InDels) are all framed in red. **B**. Picture of *Atd27like1* and *Atd27 Atd27like1* mutants in comparison with *Atd27* and Col-0 wild type. *Atd27* and *Atd27 Atd27like1* mutants have increased axillary branching. Arabidopsis plants were recorded 45 days after sowing. **C.** Plant height of Col-0, *Atd27*, *Atd27like1* and *Atd27 Atd27like1* mutants. **D**. Numbers of axillary branches in Col-0, *Atd27*, *Atd27like1* and *Atd27 Atd27like1* mutants. **E**. Striga seed germination assay conducted by applying root exudates of 5-week-old Arabidopsis plants, using GR24 (2.5 μM) as a positive control. **F.** Picture of *Atd27 AtD27LIKE1-*OX lines in comparison with *Atd27* and Col-0 wild type. Overexpression of *AtD27LIKE1* restored wild type branching to the *Atd27* mutant. **G**. Numbers of axillary branches in Col-0, *Atd27* and *Atd27 AtD27LIKE1*-OX lines.

To further investigate the biological function of *AtD27LIKE1* and to confirm its contribution to SL biosynthesis, we transformed Col-0 and *Atd27* plants with the plasmid pMDC32-*AtD27LIKE1* that enables the expression of *AtD27LIKE1* under the control of the constitutive cauliflower mosaic virus 35S promoter (CaMV 35S). We isolated three independent *Atd27 At*D27LIKE1-OX lines with more than 20-fold higher *D27LIKE1* expression levels (**Supplemental Figure S6B**). We also identified two *AtD27LIKE1*-OX lines with approximately 3-fold higher expression level of *AtD27LIKE1*, compared to the wild type (**Supplemental Figure S6A**). The number of shoot branches in the three *Atd27 AtD27LIKE1*-OX lines (Figure 3F) was similar to that of Col-0 (Figure 2G), suggesting that *AtD27LIKE1* overexpression completely restored the *d27* branching phenotype to that of the wild-type. The capability of *AtD27LIKE1* to restore *Atd27* mutant shoot branching indicates that *At*D27LIKE1 forms the SL precursor 9-*cis*-β-carotene *in planta*. To check this hypothesis, we determined the carotenoid pattern in young leaves of *Atd27*, *Atd27like1*, *Atd27 Atd27like1*, and *AtD27LIKE1*-OX lines by absolute quantification of β-carotene isomers based on the all-*trans*-β-carotene standard curve (**Supplemental Document 1**). Confirming our hypothesis, we observed around 1.3-fold increase in 9-*cis*-β-carotene content in leaves of the two *AtD27LIKE1*-OX lines (Figure 4), which could explain the observed rescue of the *Atd27* more-branching phenotype by constitutive expression of *AtD27LIKE1*. In summary, our data demonstrate that *AtD27LIKE1* contributes to the biosynthesis of SL and may play a role in determining SL homeostasis within Arabidopsis, and that this function is due to the *cis*/*cis*-isomerization activity of the encoded enzyme.

**Figure 4.**
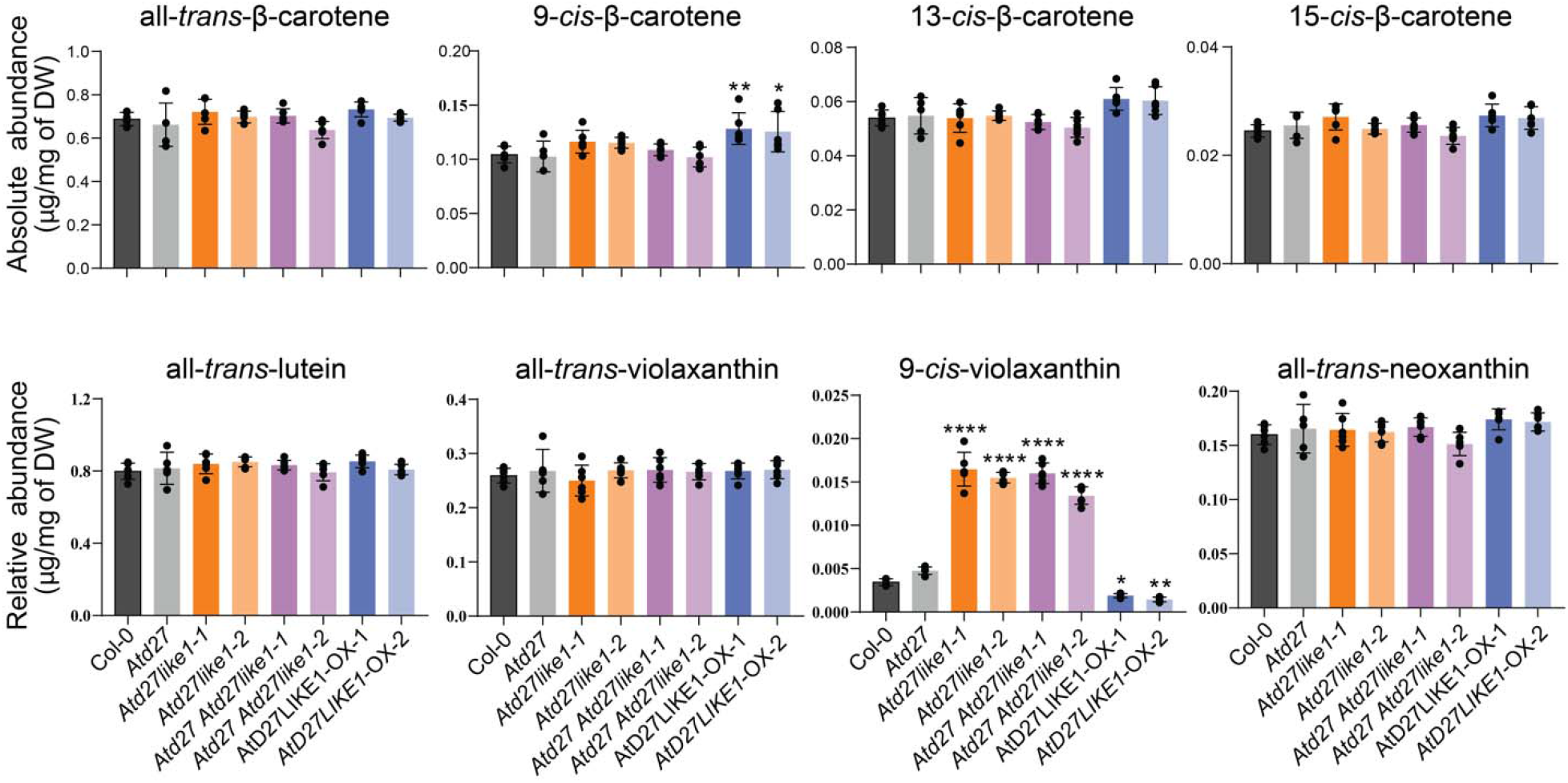
Carotenoid quantification in leaves of Arabidopsis lines with perturbed *At*D27 and/or *At*D27LIKE1 activity. Absolute abundance of all-*trans*-β-carotene, 9-*cis*-β-carotene, 13-*cis*-β-carotene and 15-*cis*-β-carotene in Col-0, *Atd27*, *Atd27like1*, *Atd27 Atd27like1* mutants and *AtD27LIKE1*-OX lines. Relative contents of all-*trans*-lutein, all-*trans*-violaxanthin, 9-*cis*-violaxanthin and all-trans-neoxanthin in Col-0, *Atd27*, *Atd27like1*, *Atd27 Atd27like1* mutants and *AtD27LIKE1*-OX lines. A two-way ANOVA with Dunnett’s test was performed to determine significance (n ≥ 5). *: p < 0.05, **: p < 0.01, ****: p < 0.0001. Error bars represent ±SD. DW: dry weight.

### Disruption of *AtD27LIKE1* increased 9-*cis*-violaxanthin and ABA content

Carotenoid profiling (**Supplemental Document 1**) of *AtD27LIKE1* knock-out and overexpressing lines unraveled a clear impact on the content of 9-*cis*-violaxanthin in leaves. As shown in Figure 4 and 5B, the disruption of *AtD27LIKE1* caused a 4-fold increase in the amount of 9-*cis*-violaxanthin, while its constitutive overexpression had an opposite effect, i.e. around 50% decline, compared to the wild type. We also quantified highly abundant xanthophylls, including all-*trans*-lutein, -violaxanthin, and -neoxanthin based on the calibrated all-*trans*-lutein standard curve (**Supplemental Document 1**); however, we did not observe any significant changes (Figure 4). Considering that 9-*cis*-violaxanthin is a precursor of ABA, we quantified the ABA level in leaves of 10-days seedlings of *AtD27LIKE1* knock-out and overexpressing lines, using the ABA deficient mutant *aba1-6* as a negative control. We detected a significantly enhanced ABA level (Figure 5A) in the Arabidopsis mutant lines disrupted in *AtD27LIKE1*, and as expected, the *aba1-6* mutant contained a very low ABA amount. We did not observe significant differences between wild type and *Atd27* and *AtD27LIKE1*-OX lines (Figure 5A). We also checked the transcript levels of *NCED* genes catalyzing the rate-limiting step in the ABA biosynthesis. Notably, the expression levels of the five *NCED2*, -*3*, -*5*, -*6*, and -*9* present in Arabidopsis (Iuchi et al., 2001) were significantly upregulated in leaves of seedlings with the *Atd27like1* background, which was consistent with the higher ABA content (**Supplemental Figure S7**). We also observed higher expression levels of *NCED2, 3,5,9* in the *Atd27* line, but to a lesser extent than in the *Atd27like1* background. These data indicated a negative correlation between *Atd27like1* presence/expression level and ABA content. Confirming the negative impact of *AtD27LIKE1* on ABA content, we observed a relatively higher ABA content in *Atd27like1* and *Atd27 Atd27like1* dry seeds (Figure 5C), while *Atd27* seeds contained less ABA. In accordance with higher ABA abduance, *Atd27 Atd27like1* mutants showed a late germination (Figure 5D) from 30h to 36h after sowing. *Atd27like1-2* displayed decreased germination ratio at 36 hour. (Figure 3E), *Ataba1-6* exhibited earlier germination ratio than that of wild-type.

**Figure 5.**
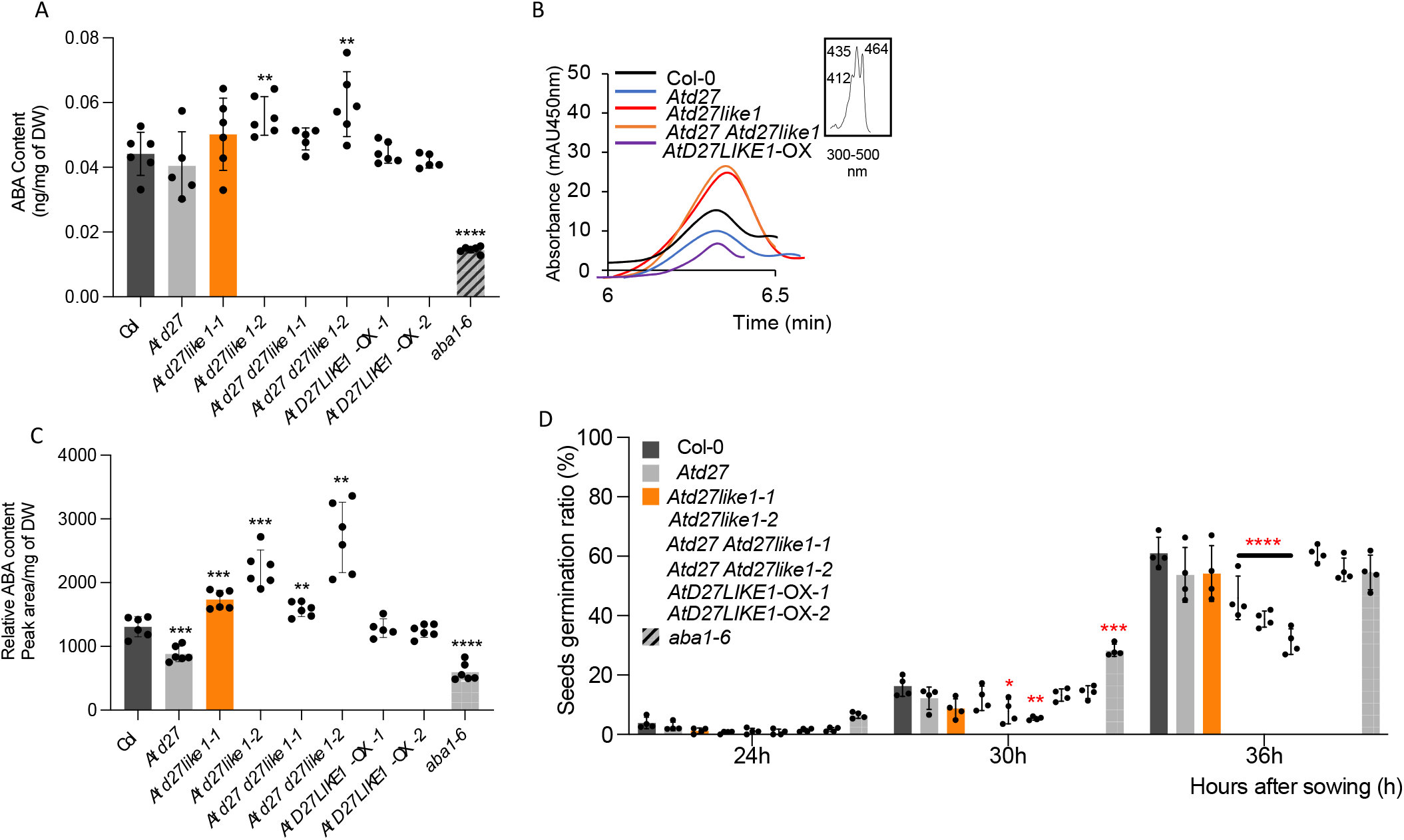
ABA content in shoots and seeds of different mutant lines and seeds germination phenotype. **A.** ABA content in shoot tissues of Col-0, *Atd27, Atd27 Atd27like1, AtD27LIKE1*-OX plants. **B.** Pattern of 9-*cis*-violaxanthin in Col-0, *Atd27, Atd27like1, Atd27 Atd27like1* mutants and *At*D27LIKE1-OX lines. UV-Vis spectra are depicted in the insets. **C.** Relative ABA content in dry seeds of Col-0, *Atd27, Atd27 Atd27like1, AtD27LIKE1-OX* lines as well as *aba1-6* as negative control. **D.** Seeds germination ratio in Col-0, *Atd27, Atd27 Atd27like1, AtD27LIKE1-*OX and *aba1-6* line. Germination ratio was recorded starting from 24h after sowing. A non-paired two-tailed Student’s *t*-test was performed to determine the statistical significance (*n* =3). *: p < 0.05, **: p < 0.01, ***: p < 0.001, ****: p < 0.0001.

### *AtD27LIKE1* is expressed in different tissues and is regulated by ABA and Auxin

To determine the expression pattern of *AtD27LIKE1* in different Arabidopsis tissues, we generated three independent Arabidopsis transgenic lines carrying the expression cassette of a nuclear localized-GUS reporter gene driven by the *AtD27LIKE1* promoter. Interestingly, GUS signal appeared in various tissues and different growth stages of T_3_ lines; for instance, *AtD27LIKE1* was predominantly expressed in young leaves (Figure 6A), anchor root primordia (Figure 6B), the differentiation zone of lateral root (Figure 6C), and stem vascular tissues (Figure 6D) of 7-day-old Arabidopsis seedlings. The *AtD27LIKE1* promoter was also active in 5-week-old Arabidopsis plants, as strong GUS signals were detected in the internodes (Figure 6E), trichomes (Figure 6F) of leaves, stigma (Figure 6G) and style (Figure 6H) of mature flowers. Furthermore, *AtD27LIKE1* expression was detected in the radicle (Figure 6I) of mature seeds. We further investigated the impact of plant hormones on *AtD27LIKE1* expression by using qRT-PCR and GUS expression assays enabled by the *pAtD27LIKE1:NLS-GUS* expression in transgenic Arabidopsis lines. Application of ABA led to an increase in the transcript levels of *AtD27LIKE1* in both shoots and roots, while 1-naphthaleneacetic acid (NAA, a synthetic auxin) treatment caused an enhancement only in root tissues. Application of the SL analog MP3 (Jamil et al., 2018) and the growth regulator zaxinone (ZAX) (Wang et al., 2019) did not affect the transcript levels of *AtD27LIKE1*(Figure 6J, L). For the *GUS*-reporter studies, we treated uniform, 14-day-old *pAtD27LIKE1:NLS-GUS* seedlings with 50 μM ABA, 1 μM NAA, 10 μM MP3, and 50 μM ZAX for six hours. The ABA treatment increased the GUS activity in the differentiation zone of lateral roots(Figure 6K), and in trichomes (Figure 6M), while NAA treatment significantly induced the GUS signal in lateral roots. Consistent with qRT-PCR data, we did not observe any effect of MP3 or zaxinone treatment (Figure 6K, M).

**Figure 6.**
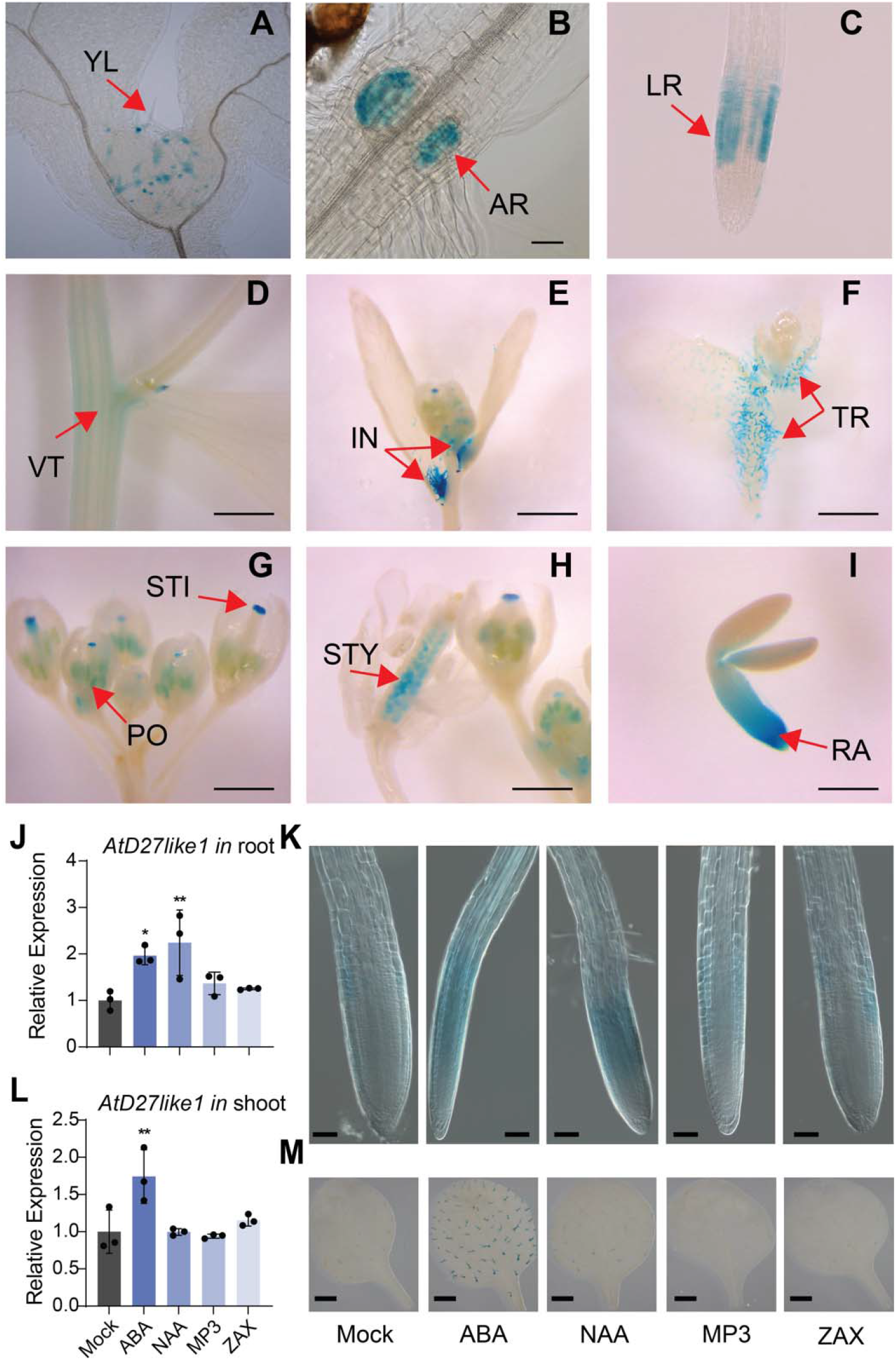
Histochemical localization of *AtD27like1*-GUS expression in transgenic *Arabidopsis* and effect of hormone treatment on *At*D27LIKE1 expression. **A.** The expression of *AtD27like1* in 7-day-old seedlings in young leaves (YL), **B**. anchor root primordia (AR), **C**. the differentiation zone of lateral root (LR), **D**. vascular tissue in the stem (VT), **E**. internodes (IN) of 5-week-old plant, **F**. trichomes (TR), **G**. stigma (STI) and pollen (PO) of the immature flowers, **H**. style (STY) and stigma (STI) of the mature flowers, **I**. radicle (RA) of mature seed. Scale bars: (A). 20 μm, (B, C, D). 50 μm, (E, F, G, H, I). 500 μm. **K**. and **M**. Expression of *AtD27LIKE1* in root tissues and shoot tissues of 14-day-old wild-type Col-0 seedlings upon treatment with 50 μM ABA, 1 μM 1-Naphthaleneacetic acid (NAA), 10 μM methyl phenlactonoate 3 (MP3, an SL analog) and 50 μM Zaxinone (ZAX). The *AtACTIN* was used as a housekeeping reference gene. A non-paired two-tailed Student’s *t*-test was performed to determine the statistical significance (*n* =3). *: p < 0.05, **: p < 0.01. Bars represent means ± SD. **J** and **L**. *pAtD27LIKE1:NLS-GUS* expression in the primordia of lateral roots (J) and trichomes of shoots (L). Scale bars: (K). 20 μm, (M). 1mm.

## DISCUSSION

D27 was initially characterized as an iron-binding protein of unknown function, which is involved in SL biosynthesis in rice (Lin et al., 2009). The strict substrate stereospecificity of CCD7 and the role of D27 acting upstream of MAX1 that is localized in the cytoplasm indicated that D27 might be the enzyme responsible for the isomerization of the predominant all-*trans*-β-carotene to 9-*cis*-β-carotene, the substrate of CCD7. *In vivo* assays and *in vitro* activity tests revealed that the rice and the Arabidopsis D27, i.e. *Os*D27 and *At*D27, catalyze the reversible isomerization of all-*trans*-β-carotene into 9-*cis*-β-carotene (Bruno and Al-Babili, 2016; Abuauf et al., 2018). D27 was the first and only β-carotene isomerase identified in plants. Sequence and phylogenetic analysis unraveled the presence of two D27 homologs, D27LIKE1 and D27LIKE2. D27 and D27LIKE1 are present in all land plants, while D27LIKE2 is also identified in some chlorophytes and prokaryotes with 3 cyanobacterial proteins referred in an outgroup, indicating a single D27-like gene copy served as the common ancestor of the chlorophyte and land plants. Phylogeny showed the first of two gene duplication events occured in the lineage, giving rise to the D27 and D27LIKE1 in the land plants, and D27LIKE2 arose earlier and exerted distinct function from D27 and D27LIKE1(Waters et al., 2012). D27 and D27LIKE1 proteins share a common ancestor origin, making it tempting to assume that D27LIKE1 might keep a similar function as D27 (Waters et al., 2012). We observed the active reversible isomerization of 9-*cis* to 13-*cis-*β-carotene, undirectional isomerization from 15-*cis*-to 13-*cis*- and 9-*cis*-β-carotene, and 9-*cis*-to all-*trans-*β-carotene, indicating the wide isomerization activity of *At*D27LIKE1 at the C_9_-C_10_, C_13_-C_14_ and C_15_-C_15’_ double bonds (Figure 2A-E). This activity was not reported before. Plants synthesize carotenoids in plastids, which derived from endosymbiotic cyanobacteria during evolution. Therefore, we tested the isomerization activity of *At*D27LIKE1 in the cyanobacterium *Synechocystis* sp. PCC 6803. Although, we did not detect a significant change in 9-*cis*-β-carotene content, the observed decrease in the amounts of other *cis*-β-carotene confirmed the results obtained *in vitro*, which identified 13-*cis*- and 15-β-carotene as substrates of AtD27LIKE1. It might be speculated that the enzyme converts 13-*cis*- and 15-*cis*-β-carotene into 9-*cis*-β-carotene and that the latter cannot be increased to levels higher than those observed in *ΔcrtO* and is degraded either directly or after being converted into all-*trans*-β-carotene that is also present at saturating concentrations.

The innovative generation of 9-*cis-*β-carotene from other *cis*-isomers catalyzed by *At*D27LIKE1 suggested its possible involvement in SL biosynthesis. For this purpose, we generated single *Atd27like1* and double *Atd27 Atd27like1* CRISPR-Cas9 knock-out mutants, overexpressed *AtD27LIKE1* in *Atd27* as well as in wild type Col-0 and performed phenotypic and metabolic analysis in all the above-mentioned lines. Arabidopsis *Atd27like1* mutants displayed a normal branching phenotype as wild type and *Striga* seed germination bioassay results showed that the SL contents in the root exudates of wild type and *Atd27like1* were at a similar level (Figure 5G). However, more severe SL-related phenotype in *Atd27 Atd27like1* lines and overexpression of *AtD27LIKE1* in *Atd27* completely restored its high-branching phenotype indicated the redundant role of *At*D27LIKE1 in SL biosynthesis, which was further proven by the increased 9-*cis-*β-carotene content in the *AtD27LIKE1*-OX lines (Figure 5G).

The presence of D27 and D27LIKE1 in the moss *Physcomitrella patens* indicated each of the two groups originated from a second gene duplication event that occured during the early evolution process of land plants (Waters et al., 2012). A novel gene function should be acquired in the evolution process during a gene duplication event as multicellularity evolved (Waters et al., 2012). Thus, we propose *At*D27LIKE1 exert different roles besides harboring redundant functions involved in SL biosynthesis. The increased ABA and 9-*cis*-violaxanthin level observed in the Arabidopsis *AtD27like1* knock-out background lines suggested that *At*D27LIKE1 might be involved in the regulation of ABA biosynthesis. In contrast to our observation, less drought tolerance and decreased ABA content in rice *OsD27like1* shoot was reported(Liu et al., 2020). The decreased endogenous ABA level in the rice *OsD27like1* as that of *OsD27* mutant might be explained from the more ancient role in regulating ABA biosynthesis in the conserved D27 family(Haider et al., 2018). The requirement of isomerization from all-*trans*-β-carotene and all-*trans*-violaxanthin/neoxanthin in SL and ABA biosynthesis is equivalent. Our results excluded the direct involvement in formation of 9/9’-*cis* counterparts for ABA biosynthesis(**Figure S3**). It seems 9-*cis*-β-carotene negatively impacts the accumulation of 9-*cis*-violaxanthin as the *AtD27LIKE1*-OX lines showed higher 9-*cis*-β-carotene but less 9-*cis*-violaxanthin (Figure 7). Both *AtD27LIKE1*-OX lines contained less 9-*cis*-violaxanthin and similar ABA levels, the latter was probably due to far-more abundant 9’-*cis*-neoxanthin was the main ABA precursor and the non-distinctive decline of 9-*cis*-violaxanthin. Although ABA levels in shoots of *Atd27like1-2* and *Atd27 Atd27like1-2* were higher than in *Atd27like1-1* and Atd27 *Atd27like1-1*, dry seeds displayed the similar trend but in which ABA contents were all significantly more than that in wild-type(Figure 5A,C). Double mutants all exhibited later germination compared to wild-type, which were consistent with higher ABA levels in dry seeds.

**Figure 7.**
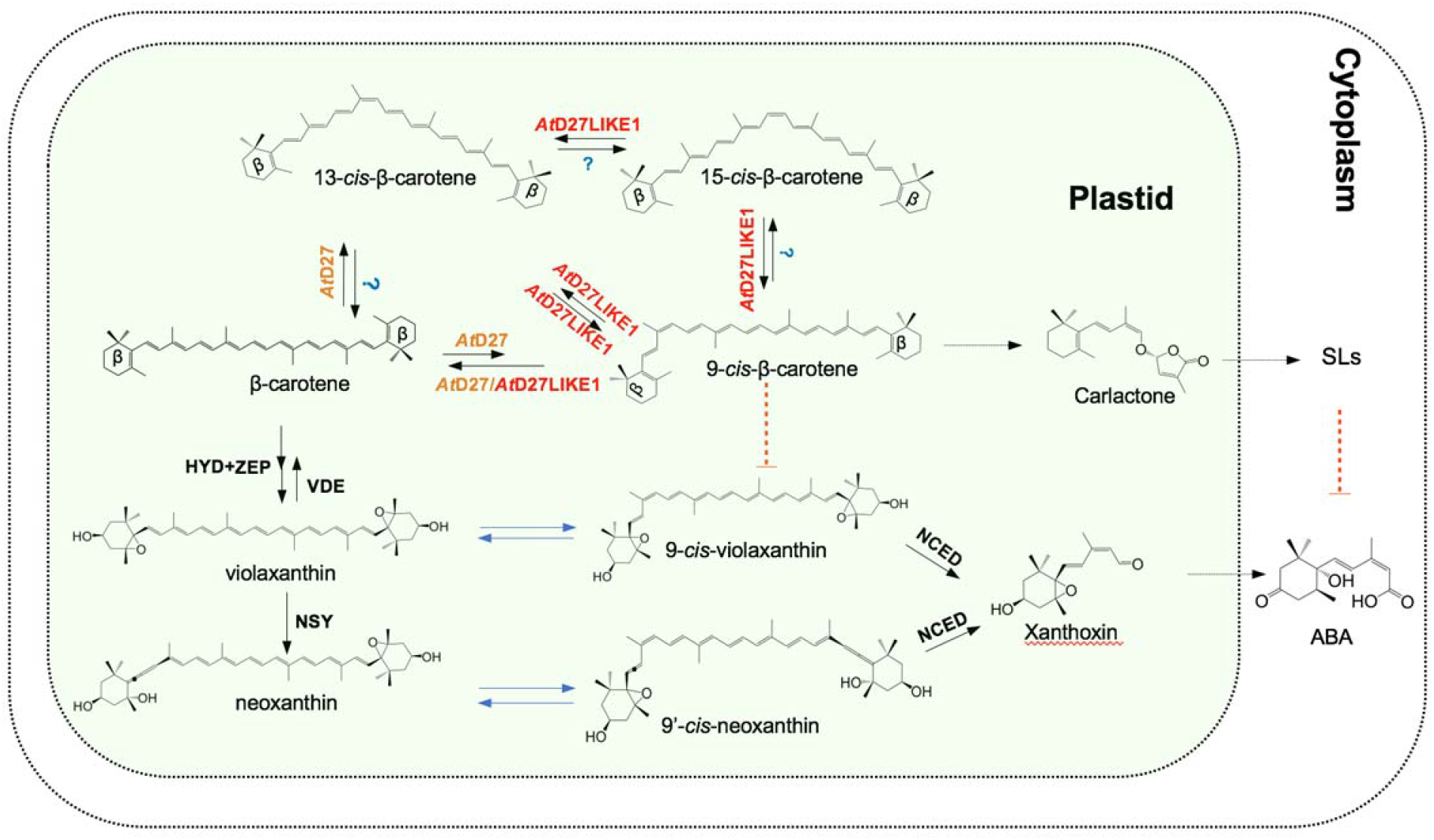
Hypothetical model for the roles of *At*D27LIKE1 in SL and ABA biosynthesis. HYD, NON-HEME DIIRON OXIDASE; ZEP, ZEAXANTHIN EPOXIDASE; NSY, NEOXANTHIN SYNTHASE; VDE, VIOLAXANTHIN DE-EPOXIDASE; NCED, 9-*CIS*-EPOXYCAROTENOID DIOXYGENASE; Unknown enzymes are indicated with blue color; the dotted arrow indicates multiple reactions.

SLs and ABA are all carotenoid-derived phytohormones, it woud be of interest to study how the levels of SL and ABA are fine-tuned during growth and development. Disruption of ABA biosynthesis leads to the also down-expression of SL biosynthesis genes in tomato and Arabidopsis (López-Ráez et al., 2010; Van Ha et al., 2014). However, elevated expression of *D27, D20* and D17 was observed in ABA-deficient rice, and ABA treatment led to the reduction of SLs in wild-type. All indicated the inhibitory effect of ABA on SL biosynthesis in rice and the disctinct regulation according to species specificity (Liu et al., 2020). SLs were reported to induce the expression of *AtHB40*, which encoded a BRC1-target transcription factor that promoted ABA biosynthesis through activating transcription of *AtNCED3*(Wang et al., 2020). However, on the contrary to the above-findings, our observations show it’s likely SLs negatively influences the metabolism of ABA through the flux regulation of 9-*cis*-violaxanthin from 9-*cis*-β-carotene(Figure 7), which is connected through *AtD27LIKE1* with unknown mechanism. Perhaps, the difference might be somehow resulted from the complicated and indirect connection between metabolites and gene expression. It was observed that induced expression of ABA biosynthesis gene(β-carotene hydroxylase, *BCH*) leads to the accumulation of ABA-pathway intermediates rather than the increased ABA levels(Du et al., 2010).

In summary, we have characterized *At*D27LIKE1 and shown that it catalyzes an isomerization activity converting *cis*-isomers into each other, which increases 9-*cis*-β-carotene content *in planta*. Our data indicate that *At*D27LIKE1 contributes to SL biosynthesis, while it negatively impacts ABA content in shoots and seeds (Figure 5), which adds another connection layer between the biosynthetic pathways of ABA and SLs.

## EXPERIMENTAL PROCEDURES

### Carotenoid substrates used in *in vivo*/*in vitro* assays

Synthetic carotenoids, including all-*trans*, 9,13, 15-*cis*-β-carotene, all-*trans*-α-carotene, all-*trans*-violaxanthin, all-*trans*-neoxanthin were purchased from CaroteNature (Lupsingen, Switzerland). Lutein and 9’-*cis*-neoxanthin were extracted and purified from fresh spinach obtained from the local supermarket (Altemimi et al., 2015). Carotenoid substrates were quantified photometrically based on their molar extinction coefficients(Britton, 1995). All the reagents used in this work were of the highest possible quality.

### *In vitro* assays with Arabidopsis D27 and D27LIKE1 enzymes

BL21(DE3) *E.coli* competent cells transformed with pGro7 (Takara Bio Inc), harboring the chaperons that assist in the folding and assembly of target proteins, were transformed with the empty plasmid pThio-Dan1 (Trautmann et al., 2013), pThio-Dan1-*At*D27(-cTP), and pThio-Dan1-*At*D27LIKE1(-cTP). Positive clones were inoculated into 5 mL 2YT medium (containing 1.6% (w/v) tryptone, 0.2% (w/v) glucose, 0.5% (w/v) NaCl and 1% (w/v) yeast extract) and grown at 37°C overnight. 1 mL of overnight culture was inoculated into 50 mL 2YT medium and incubated at 37°C until an OD_600_ of 0.5. Then, 0.2% arabinose (w/v) was added to induce the protein expression, and cultures were incubated at 28°C for another 4 h. Cells were centrifuged at 1600 g for 15 min, and pellets were resuspended with 1 mL lysis buffer containing 0.1 M PBS (Phosphate-buffered saline), 1mM DTT (Dithiothreitol) and 0.1% TritonX-100, 10 mg/mL lysozyme and incubated on ice for 30 min, followed by sonication (3 times each 5 sec and centrifugation at 21,000 g at 4°C for 10 min. The supernatant carrying the crude lysate fraction was used as an enzyme preparation to perform *in vitro* assays. *In vitro* assays were performed in a total volume of 200 µL with 100 µL assay buffer composed of 0.22 mM FeSO_4_, 2mM TCEP(tris(2-carboxyethyl)phosphine), 200 mM HEPES ((4-(2-hydroxyethyl)-1-piperazineethanesulfonic acid) (pH=8), 2 mg/mL catalase, 50 µL crude soluble enzyme and 60 μM substrate. The assays were performed at 28°C for 30 min under agitation at 200 rpm in darkness. Extraction was performed with two volumes of acetone, followed by sonication (3 times each 5sec, and then two volumes of petroleum/diethyl ether (1:4, v/v) were added. After vortex and short centrifugation (⁓10 sec), the organic phase was collected and dried in a vacuum centrifuge. The dried samples were re-dissolved in 80 µL CHCL_3_ and filtered with 0.2 μM filters, and 10 µL of the samples were subjected to UHPLC analysis as explained in the UHPLC analysis section.

### *In vivo* assays with Arabidopsis D27 and D27LIKE1 enzymes

The empty vector pThio-Dan1 (Trautmann et al., 2013), pThio-Dan1-*At*D27(-cTP), and pThio-Dan1-*At*D27LIKE1(-cTP), were transformed into transgenic *E. coli* strains that accumulate β-carotene, lycopene and zeaxanthin (Matthews and Wurtzel, 2000; Prado-Cabrero et al., 2007). Positive clones were selected and grown overnight at 37°C in 5 mL Luria-Bertani (LB) medium containing 100 mg/L ampicillin and 50 mg/L kanamycin. Then, 1 mL of overnight culture was inoculated into 50 mL of LB medium and incubated at 37°C until reaching an OD_600_ value of 0.5. Each thioredoxin-corresponding protein expression (*At*D27 or *At*D27LIKE1) was induced by 0.2% of arabinose (w/v), and cultures were further incubated for 4 h at 28°C covered with aluminum foil to prevent isomerization by light. Cultures were harvested by centrifugation at 1600 g for 15 min, and 2 mL of acetone was added to the harvested pellets, followed by sonication (3 times each 5 s). After centrifugation for 10 min at 1600 g, the supernatants containing organic compounds were dried in a vacuum centrifuge. Organic samples were dissolved in 1 mL chloroform (CHCL_3_) and dried again in a vacuum centrifuge. Dried samples were re-dissolved in 80 µL CHCL_3_ and filtered through 0.2 μM filters with 10 µL, and then subjected to UHPLC analysis with system 1, system 2, and system 3, respectively.

### UHPLC analysis

UHPLC ultimate 3000 system (Thermo Fisher Scientific) equipped with UV detector and autosampler was used for carotenoids analysis. The LC separations of β-carotene and lycopene were carried out with the YMC-pack-C_30_ reversed column (150×30 mm, 5μm, YMC Europa, Schermbeck, Deutschland) kept at 30°C with mobile phase A (methanol: tert-butylmethyl ether (1:1, (v/v) and mobile phase B (methanol: water: tert-butylmethyl ether (5:1:1, (v/v/v)). Eluting gradient system for β-carotene (System 1) started from equilibration with 80% B for 8 min, increased to 100% A for 20 min, followed by washing with 100% A for 5 min. The separation system for lycopene (System 2) was as follows: 0-8 min 70% A, 8-28 min, 70%-100% A, followed by washing with 100 % A for 5 min. The chromatographic separation of xanthophylls, α-carotene, and carotenoids were performed on a YMC-pack-C_18_ reversed column (150×30 mm, 3μm, YMC Europa, Schermbeck, Deutschland) maintained at 30°C. Mobile phases used for xanthophylls and α-carotene were A (methanol: tert-butylmethyl ether (1:1, (v/v) and B (methanol: water: tert-butylmethyl ether (30:10:1, (v/v/v)). The eluting program (System 3) for xanthophylls and α-carotene started from equilibration with 30% A for 8 min, with a gradient of A increasing from 30% to 100% within 24 minutes, maintaining the final conditions for 1 min. The chromatographic separation of total carotenoids (System 4) was performed with a gradient from 30% A to 100% A (A, methanol: tert-butylmethyl ether (1:1, v/v; and B, methanol: water: tert-butylmethyl ether (6:3:1, (v/v/v)) in 19 min, maintaining 100% A for 5 min, decreasing 100% A to 70% A in 1 min and washing of 5 min with 70% A. The flow rate was 0.6 ml/min in enzymatic assays and 0.8 ml/min in the total carotenoids separation system.

### Plant materials and growth conditions

Wild-type Arabidopsis (Col-0), *aba1-6* (CS3772) (Barrero et al., 2005), *Atd27* T-DNA insertion line with Col-0 background (Waters et al., 2012) and all transgenic lines were sterilized and stratified at 4°C in the dark for 3 days. For phenotypic characterization, seeds were sown on ½ MS agar (0.5 MS, 1% sucrose, 1% agar, 0.5 g/L 2-(N-morpholino) ethanesulfonic acid, pH 5.7) plates and kept in Percival growth chamber (Biochambers) under long-day photoperiod (22°C/20°C 16 h light/8 h dark; 150 µmol m^−2^ s^−1^) for 14 days. Seedlings were transferred to the plant growth facility (16 h light/8 h dark; 22°C/20°C; 120 µmol m^−2^ s^−1^) for 31 days. For carotenoids analysis, sterilized seeds were sown on ½ MS-agar plates and grown in the same chamber for 10 days. For GUS-staining, seeds were grown on ½ MS-agar plates for 7 days. Another batch of seeds was grown on agar plates for 10 days and transferred into soil, and then grown in the growth facility for 25 days.

For Striga seed germination bioassays, sterilized and stratified seeds were grown in a hydroponic system adopted from (Conn et al., 2013) with a modified half-strength Hoagland nutrient solution (0.4mM K_2_HPO_4_⋅3H_2_O, 0.8 mM MgSO_4_⋅7H_2_O, 0.18 mM FeSO_4_⋅7H_2_O, 5.6 mM NH_4_NO_3_, 0.8 mM K_2_SO_4_, 0.18 mM Na_2_EDTA⋅2H_2_O; and 23 μM H_3_BO_3_, 4.5 μM MnCl_2_⋅4H_2_O, 1.5 μM ZnCl_2_, 0.3 μM CuSO_4_⋅5H_2_O, and 0.1 μM Na_2_MoO_4_⋅2H_2_O) with an adjusted pH value of 5.75 in the Percival growth chamber (Biochambers) under (22°C, 10 h light/14 h dark, 100 µmol m^-2^ s ^-1^ light intensity, 60% humidity). The nutrient solution was replaced twice a week. After four weeks, the plants were transferred into the 50 ml black tubes and grown for another week with the replacement of a phosphate (K_2_HPO_4_.2H_2_O) free nutrient solution.

For seed germination assay, seeds including Col-0, *Atd27*, *Atd27like1*, *Atd27 Atd27like1* and *AtD27LIKE1*-OX lines as well as *Ataba1-6* as a negative control were sown on ½ MS agar (without sucrose) plates and grown in the same chamber and radicle emergence was recorded as germination at different time points. In accordance with a previously published report, the time required for Arabidopsis wild-type Col-0 to reach radicle emergence, hypocotyl and cotyledon emergence was 1.3- and 2.5-day, respectively (Boyes et al., 2001)

For hormone and apocarotenoid treatments, *pAtD27like1:NLS-GUS* and Col-0 seedlings were grown on ½ MS agar plates and kept in a Percival growth chamber (Biochambers) under long-day photoperiod (22°C/20°C 16 h light/8 h dark; 150 µmol m^−2^ s^−1^) for 14 days (Abuauf et al., 2018). Uniform seedlings were selected and transferred to ½ Hoagland nutrient solution for 6 h treatment with the indicated concentrations of chemicals (Mock: no addition, 1 μM synthetic auxin 1-naphthaleneacetic acid (NAA), 50 μM ABA, 10 μM synthetic SL analog methyl phenlactonoate (MP3) (Jamil et al., 2018) and 50 μM zaxinone (Wang et al., 2019). The *pAtD27like1:NLS-GUS* seedlings were conducted with GUS staining according to a previously described protocol (Jefferson et al., 1987), and the GUS signals were examined and photographed from stained roots and leaves.

### Plasmid construction, mutant isolation, and generation of transgenic plants

The pThio-Dan1-*At*D27 was reported in (Abuauf et al., 2018). The pThio-Dan1-*At*D27LIKE1 and *pAtD27like1:NLS-GUS* were generated according to (Abuauf et al., 2018), synthetic *AtD27like1* cDNA after removal of cTP is obtained from Genewiz (**Supplemental Document 1**). The single *Atd27like1* and *Atd27 Atd27like1* double mutants were generated using CRISPR/Cas9 technology. Briefly, an *Atd27like1*-specific target sequence was designed online at http://www.e-crisp.org/E-CRISP/designcrispr.html. Single-guide RNA (sgRNA) cassette was cloned into the pHEE401 vector (Wang et al., 2015). To produce transgenic plants overexpressing the *Atd27like1* gene, full-length cDNA was subcloned into the modified binary pMDC32 vector under the control of the CaMV 35S promoter. The constructs were transformed into Arabidopsis Col-0 and *Atd27* mutant plants by the floral dip method (Clough and Bent, 1998). The resulting plants were selected on MS medium containing hygromycin (50 mg/L). To retain overexpressing lines that are presumed to contain a single copy of the transgene, T_1_ seeds were sown on selective antibiotic ½ MS medium (50mg/L hygromycin).The single copy T_1_ lines were determined by segregation ratio of 3:1 for selective antibiotic resistance/non-resistance. The seeds of single copy T_2_ lines were again germinated in selective antibiotic ½ MS medium, the T_2_ lines with 100% seed germination were selected for the development of generation T_3_ homozygous plants. Three independent T_3_ homozygous lines containing a single copy of the transgene were selected for further phenotyping analysis. Mutations in *Atd27like1* were confirmed by Sanger sequencing, and overexpression lines were confirmed by qRT-PCR.

### Generation of transgenic *Synechocystis* strains

A *Synechocystis* codon-optimised version of *AtD27like1* (At1g64680) was (synthesized by Genewiz, USA; see **Supplemental Table S1**) fused with a 6 X His-tag sequence on the C-terminal. Subsequently, *AtD27like1* was constructed under the control of a constitutive promoter Pcpc and terminator of TrpnB using CyanoGate Kit (#1000000146, Addgene, USA). To insert the *AtD27like1* gene cassette in the *crtO* gene site, around 1000 bp upstream and downstream flanking regions of the *crtO* gene were amplified as homology arms by Phusion high-fidelity DNA polymerase (#M0530L, NEB) from *Synechocystis* genomic DNA using specific primers (**Supplemental Table S1**) and cloned into pJET 1.2 (CloneJET PCR Cloning Kit # K1231, Thermo Fisher Scientific). Finally, the gene cassette, homology arms for integration and kanamycin-resistance cassette were combined and ligated into the acceptor vector pCAT.334. The construct was transformed into wild-type *Synechocystis,* the resulting strains were selected on solid BG-11 plates containing kanamycin (50 mg/L), and expression of *AtD27like1* was confirmed by PCR and Western-Blot analysis.

### *Synechocystis* Strains and culture conditions

The glucose tolerant *Synechocystis* sp. PCC 6803 strain was used as a wild type. The mutant Δ*crtO* with deletion of the gene *crtO* was generated in our lab. All *Synechocystis* strains were cultured at 30°C under constant illumination (50 μmol m ^−2^ s ^−1^) in BG-11 medium (Rippka et al., 1979) supplemented with appropriate antibiotics. For liquid culture, cells densities were monitored by measuring the absorbance at 750 nm using a spectrophotometer (Thermo Fisher Scientific). For sample collection, all cultures were diluted to 0.2 (OD_750_) and grown for two weeks in sterilized flasks. Three independent biological replicates were performed. Solid BG-11 plates were prepared with 0.9 % Kobe agar.

### RNA isolation and qRT-PCR analysis

Total RNA was extracted from Arabidopsis seedlings using the Direct-zol RNA MiniPrep Plus Kit (Zymo Research). Briefly, 1 μg RNA was reverse-transcribed to cDNA with the iScript^TM^ cDNA synthesis Kit (Bio-Rad). Amplification with primers (Supplemental Document1) was done with the SYBR Green Real-Time PCR Master Mix kit (Life Technologies). The qRT-PCR program was run in a StepOne Real-Time PCR System (Life Technologies). The thermal profile was 50°C for 2min, 95°C for 10min, followed by 40 cycles of 95°C for 20s, 60°C for 20s and 72°C for 20s. The quantitative expression level of *AtD27like1* was normalized to that of the housekeeping gene (*AtActin*), and relative expression was calculated according to the 2^-ΔΔCT^ method (Yang et al., 2014).

### Protein isolation from *Synechocystis* and Western Blot analysis

Total protein was extracted from *Synechocystis* wild type and transgenic cells. Then, 20 ml of cells were grown to an optical density of 1 (OD_750nm_). Cells were collected by centrifugation at 4,000 g for 10 min, immediately frozen in liquid N_2_ and stored at −80 °C for further protein extraction. The frozen cells were resuspended with 200 μl homogenization buffer (75 mM Tris-HCl, 1.5 mM EDTA(Ethylenediaminetetraacetic acid), pH 7.5) and 0.5 mm glass beads. Samples were subsequently mixed by vortex for 1 min, then frozen in liquid nitrogen for 20 s and melted under ambient air. These steps were repeated for five cycles to disrupt cells thoroughly. After centrifugation at 2,300 g for 5 min, the supernatants were collected as total protein extracts. Then, 10 μl of total protein lysate was mixed with loading buffer and boiled at 90 °C for 10 min. Samples were then loaded and separated by SDS-PAGE and electrotransferred to a nitrocellulose membrane. Western Blot experiments were conducted according to previously published protocols (LAEMMLI, 1970; Towbin et al., 1979). The blotted membrane was incubated with the commercial anti-His HRP conjugate antibody kit (Qiagen). The signal was probed using Pierce^TM^ ECL western blotting detection reagent (#32106, Thermo Fisher Scientific). The blotting image was analyzed with the ChemiDoc XRS+ imager (Bio-Rad).

### Carotenoid extraction and quantification in *Synechocystis* sp. PCC6803 and plants

Arabidopsis green tissues were lyophilized overnight and grounded. Around 10 mg of dry weight tissue were weighted for carotenoid extraction. Initially, 1 mL acetone with 0.1% BHT(butylated hydroxytoluene) and 2-3 steel balls were added. The solution was under agitation at 300 rpm and at 4°C for 30 min and then centrifuged for 10 min at 1485 g. For carotenoid extraction from *Synechocystis,* 5 mL liquid culture was used, 2 mL acetone with 0.1% BHT was added and mixed thoroughly, the solution was sonicated in the water bath for 10 min and then centrifuged for 8 min at 1600 g at 4°C. The supernatant was transferred to a new 8 mL brown glass tube, and extraction was repeated with the pellets. Combined supernatants were dried in the vacuum centrifuge. Dried samples were dissolved in 100 µL acetone, filtered through 0.2 µm filters, and 10 µL of the samples were subjected to UHPLC analysis with system 4.

### Measurement of total chlorophyll-a and carotenoids in *Synechocystis*

To extract pigments from *Synechocystis* cells, 2 mL of cell culture with an optical density of 0.8 (OD_750nm_) was harvested by centrifugation at 4000 g for 7 min at 4 °C. The following process was performed under dim light to avoid pigment degradation. Pellets were resuspended with pre-cooled methanol and thoroughly mixed by vortex for 1 min. Samples were subsequently covered with aluminum foil and incubated at 4°C for 20 min. Cell debris was removed by centrifugation at 15,000 g for 7 min at 4 °C. The color of pellets should be purple, indicating complete pigment extraction. Methanol was used as blank, the lysate was measured at 470nm, 665nm and 720nm by spectrophotometer for concentration calculation. Three independent biological and technical replicates were performed for data analysis. Chlorophyll-a and carotenoid concentrations were calculated according to the following equations:

Chla (μg/mL) = 12.9447 (A_665_ − A_720_) (Ritchie, 2006);

Carotenoids (μg/mL) = [1,000 (A_470_ − A_720_) − 2.86 (Chla [μg/mL])]/221 (Wellburn, 1994).

### Quantification of ABA using UHPLC-MS

ABA extraction from plants was performed according to a previously published protocol (Wang et al., 2021). Briefly, ∼10 mg of freeze-dried tissue was spiked with internal standards D_6_-ABA (2 ng) with 1.5 mL of methanol. The mixture was sonicated for 15 min in an ultrasonic bath (Branson 3510 ultrasonic bath), followed by centrifugation for 10 min at 14,000 g at 4°C. The supernatant was collected with the pellet and re-extracted with 1.5 mL of the same solvent. Combined supernatants were dried under a vacuum. The dried samples were re-dissolved in 150 μL of acetonitrile:water mixture (25:75, v/v) and filtered through a 0.22 μm filter for further LC-MS analysis. ABA quantification was performed using an HPLC-Q-Trap-MS/MS (QTRAP5500; AB Sciex) and UHPLC-Triple-Stage Quadrupole Mass Spectrometer (Thermo ScientificTM AltisTM) with Multiple Reaction Monitoring (MRM) mode. Chromatographic separation was achieved on a ZORBAX Eclipse plus C_18_ column (150 × 2.1 mm; 3.5 μm; Agilent). Mobile phases consisted of water:acetonitrile (95:5, v/v) and acetonitrile, both containing 0.1% formic acid. A linear gradient was optimized as follows (flow rate, 0.4 mL/min): 0-17 min, 10% to 100% B, followed by washing with 100% B and equilibration with 10% B. The injection volume was 10 μL, and the column temperature was maintained at 40°C for each run. Mass spectrometry was conducted in electrospray and MRM mode in negative ion mode. Relevant instrumental parameters of QTRAP5500 were set as follows: ion source of turbo spray, ion spray voltage of (-) 4500 V, curtain gas of 25 psi, collision gas of medium, gas 1 of 45 psi, gas 2 of 30 psi, turbo gas temperature of 500°C, entrance potential of −10 V. The MS parameters of Thermo ScientificTM AltisTM were as follows: negative ion mode, ion source of H-ESI, ion spray voltage of 4500 V, sheath gas of 40 arbitrary units, aux gas of 15 arbitrary units, sweep gas of 10 arbitrary units, ion transfer tube gas temperature of 350 °C, vaporizer temperature of 350 °C, collision energy of 17 eV, CID gas of 2 mTorr, and full width at half maximum (FWHM) 0.4 Da of Q1/Q3 mass. The characteristic MRM transitions (precursor ion → product ion) were 263.2→153.1; 263.2→204.1 for ABA; 269.1→159.1; 269.1→207 for D_6_-ABA.

### GUS staining

The *pAtD27like1:NLS-GUS* plasmid was transformed into *Arabidopsis* Col-0 plants via *Agrobacterium tumefaciens* (strain GV3101). More than three independent T_2_ transformation lines were tested for GUS (β-glucuronidase) staining, and a representative homozygous line with a single transgene was chosen for subsequent analysis. GUS staining was performed according to (Jefferson et al., 1987), and plant tissues were examined on the microscope (Axioplan Observer.Z1, Carl Zeiss GmbH, Germany) with a digital camera (Axio Cam MRC, Carl Zeiss Microimaging GmbH, Göttingen, Germany).

### Striga seed germination bioassay

Striga seeds germination bioassays were carried out as described previously (Jamil et al., 2012)

## Author Contributions

Y.Y., A.B. performed *E.coli* enzymatic assays. Y.Y., S..S. performed *Synechocystis* sp. PCC6803 assays. S.A. and H.A. generated Arabidopsis mutants and overexpression lines. Y.Y. and X..Z performed the genotyping assay. Y.Y., H.A., J.C.M. phenotyped the Arabidopsis mutant and overexpression lines. Y.Y. and J.M. performed the carotenoids analysis. Y.Y. and J.Y.W. performed the ABA quantification assay. Y.Y., A.A., and M.J. performed the Striga seed germination assays. Y.Y., H.A., and I.B performed the GUS-staining assays. Y.Y., H.A. and A.A. performed qRT-PCR assays. Y.Y. performed the Arabidopsis seeds germination assay. Y.Y., Y.A., S..S. prepared the figures and table. Y.Y., S..S,. Y.A., and J.C. M. wrote the manuscript. J.Y.W. and S.A-B. substantially revised the manscriupt. All authors read and approved the final version of the manuscript.

## Acknowledgments

This work was supported by by Baseline funding and the Competitive Research Grants 7 and 9 (CRG7, 9) given from King Abdullah University of Science and Technology to S.A-B. We thank all members of the BioActives Lab at KAUST for their helpful discussions. We also thank the members of KAUST Greenhouse core lab for their kindly support.

## Supplemental Data

**Supplemental Figure S1.** UHPLC analysis of β-carotene, lycopene, zeaxanthin accumulating *E. coli* cells expressing thioredoxin-*At*D27 (*At*D27), thioredoxin *-At*D27LIKE1(*At*D27LIKE1), or thioredoxin (Control).

**Supplemental Figure S2.** The UHPLC chromatograms of *At*D27 and *At*D27LIKE1 enzymatic activity with xanthophylls and all-*trans*-α-carotene *in vitro*.

**Supplemental Figure S3.** Carotenoid biosynthesis pathway in *Synechocystis* sp. PCC 6803.

**Supplemental Figure S4.** The UHPLC chromatograms of carotenoid profiling in Wild-type and *ΔcrtO* mutant *Synechocystis* sp. PCC 6803.

**Supplemental Figure S5.** Pigment quantification in WT, *ΔcrtO* mutant and *AtD27L1*-OX lines.

**Supplemental Figure S6.** Generation and characterization of *AtD27LIKE1*-OX and *Atd27 AtD27LIKE1-OX* lines.

**Supplemental Figure S7.** Expression levels of *NCED2,3,5,6,9* genes in shoot tissues of Col-0, *Atd27, Atd27like1*, *Atd27 Atd27like1*.

**Supplemental Document S1** Synthesized sequences. Calibrated all-*trans*-β-carotene/lutein standard curves. Arabidopsis carotenoids profiling chromatogram and primers used in this study.

**Supplemental Table S1** Primers used in this study.

**Supplemental Table S2-S5** The proportion of different β-carotene isomers of *in vitro* assays performed with crude lysates of BL21 *E. coli* cells expressing thioredoxin-*At*D27 (AtD27), thioredoxin*-At*D27LIKE1 or thioredoxin (Control) when corresponding all-*trans*-β-carotene, 9-*cis*-β-carotene, 13-*cis*-β-carotene,15-*cis*-β-carotene were used as the substrates. Data were presented as means ± SD (n=3).

